# Comorbidity structure as an inductive bias: Comparing output-head designs for multi-label prediction of diabetes and myocardial infarction complications

**DOI:** 10.64898/2026.06.18.733068

**Authors:** Wilfred Ayine Asumboya, Perfect Kwasi Agbenorhevi, Cletus Fiifi Adams, Dennis Ayinsune Ayariga, Thaddeus Adjadeh, Shamsudini Adam Ziblim, Samuel Kojo Kwofie

**Affiliations:** Department of Biomedical Engineering, University of Ghana, Accra, Ghana; Ho Teaching Hospital, Ho, Ghana

## Abstract

**Background:** Clinical complications are often predicted with separate sigmoid outputs, even when the target labels arise from related pathophysiological processes. This paper asks whether output-layer choice should reflect both predictive convenience and the biological structure assumed among complications. The central premise is that label-dependence mechanisms are explicit hypotheses about comorbidity, not generic modelling additions.

**Methods:** Output-head assumptions were compared across two clinically distinct multi-label prediction tasks. In Type 2 diabetes (T2D), six heads were evaluated for nephropathy, neuropathy, and retinopathy: independent baseline, linear additive, multiplicative, symmetric conditional random field (CRF), residual multilayer perceptron (MLP), and combined additive–multiplicative. In myocardial infarction (MI), four heads were evaluated for ventricular tachycardia, ventricular fibrillation, and atrioventricular block: independent baseline, linear additive, multiplicative, and symmetric CRF. All experiments used five training data fractions and seven independent seeds, with the same shared-backbone protocol within each disease setting.

**Results:** In T2D, the symmetric CRF gave the most consistent improvement pattern, ranking highest at full data and at the two lowest data fractions while adding only three interaction parameters. At 20% training data, it was the only interaction head whose aggregate mean exceeded the independent baseline. The residual MLP, despite 123 interaction parameters, remained below the baseline across all T2D fractions. In MI, rankings changed across fractions: the multiplicative head led at 80% and 60%, the CRF led at 100% and 20%, and the baseline led at 40%. The combined additive–multiplicative head did not improve robustness in T2D and showed the largest negative baseline-relative deviations at lower fractions.

**Conclusion:** The findings support a biology-guided view of output-layer design. A small constrained mechanism was most useful when its symmetry matched the shared microvascular structure of T2D, whereas the heterogeneous electrophysiology of MI produced no stable winner. Output-layer choice should therefore be reported and defended as an assumption about disease structure instead of a routine hyperparameter decision.

**Author summary:** Many clinical prediction models treat complications as separate outcomes, even when clinicians know they often arise together. We studied whether the last layer of a model should reflect that biological knowledge. We compared several output heads across two disease settings: Type 2 diabetes, where nephropathy, neuropathy, and retinopathy share a common microvascular origin, and myocardial infarction, where electrical complications arise from a mixture of shared and location-specific mechanisms. We found that a small symmetric CRF head was most useful in the diabetes task, especially when training data were limited, while no single interaction head dominated in myocardial infarction. This suggests that modelling comorbidity is not only a technical choice; it is a statement about how disease processes relate to one another. Our results encourage researchers to report and justify output-layer design as part of the clinical modelling argument, rather than treating it as a routine hyperparameter.

## Introduction

### The Problem

Clinical machine learning (ML) has spent the better part of a decade optimising for predictive performance. Benchmark after benchmark has been established, model after model has pushed area under the receiver operating characteristic curve (AUROC) higher, and the field has treated this trajectory as progress [1, 2]. The structural assumptions embedded in the models producing those numbers have attracted almost no scrutiny.

In multi-label clinical prediction — where a patient presents with several related complications simultaneously — the dominant modelling approach attaches independent sigmoid outputs to a shared backbone and optimises each label separately under binary cross-entropy loss [3–5]. This is computationally convenient, easy to implement, and well-supported by standard deep learning tooling. It is also a biological claim.

Predicting diabetic nephropathy and diabetic neuropathy with independent output heads asserts that chronic hyperglycaemia damages the kidney and the peripheral nerve through separate, unrelated processes—that the two complications co-occur in patients only by coincidence, and that knowing a patient is developing one tells you nothing about whether they are developing the other. A diabetologist would reject that assertion. A cardiologist would reject the same claim applied to ventricular tachycardia and ventricular fibrillation. The biology of these diseases does not produce independent complications, and a model that assumes it does, is making a claim the clinician cannot endorse.

Label independence was inherited from the multi-label classification literature as a tractable default [3, 4, 6]. That default is architectural: it operates at the output layer before any data is seen, and cannot be corrected with more training data.

Closing the gap between architectural convenience and biological plausibility requires treating every output-layer design decision as a biological hypothesis. Where the structural assumption is accurate, a model built on plausible comorbidity structure may capture biological signal more efficiently than less constrained alternatives—a possibility evaluated across every data fraction in both disease settings (Table 3 and Table 5). What that gain depends on is the central question of this paper.

### Comorbidity as a biological spectrum

Comorbidity can be understood as a biological spectrum: in some diseases the structure is clean and recoverable by a constrained architectural assumption, whereas in others it is heterogeneous enough that no single assumption may suffice and the absence of a clear winner is itself informative.

In Type 2 diabetes (T2D), chronic hyperglycaemia damages the microvasculature systemically and continuously. The same elevated glucose that scars the glomerular capillaries in the kidney simultaneously damages the endoneurial vessels supplying peripheral nerves and the pericytes lining retinal capillaries [7–12]. The generating cause is singular, operates uniformly across every patient, and acts on all three microvascular targets at once. The co-occurrence that results is stable, symmetric, and population-level. In this dataset, nephropathy and neuropathy co-occur in 21.8% of patients, nephropathy and retinopathy in 24.7%, and neuropathy and retinopathy in 24.5% (Table 1). When nearly one in four diabetic patients develops two specific microvascular complications simultaneously, the pattern points directly at the shared generating process. None of the three complications causes the others; all three are expressions of the same underlying microvascular damage. The co-occurrence structure is undirected because the biology is undirected.

**Table 1.** Dataset characteristics and pairwise complication co-occurrence. In the T2D setting, pairwise co-occurrence rates span 21.8% to 24.7%—a 2.9 percentage point range across all three pairs, consistent with a symmetric upstream mechanism. In the MI setting, the VT–VF pair co-occurs at 5.13 times the rate expected by chance, approximately twice the lift of the VF–AVB pair at 2.52, consistent with the shared ischaemic border zone substrate that links the ventricular pair and distinguishes it from atrioventricular block.

| Panel A: Type 2 Diabetes Microvascular Complications |  |  | Panel B: Myocardial Infarction Electrical Complications |  |  |  |
| --- | --- | --- | --- | --- | --- | --- |
| Complication | Positive Cases | Prevalence | Complication | Positive Cases | Prevalence |  |
| Nephropathy | 1,468 | 47.8% | Ventricular Tachycardia | 42 | 2.5% |  |
| Neuropathy | 1,492 | 48.6% | Ventricular Fibrillation | 71 | 4.2% |  |
| Retinopathy | 1,547 | 50.4% | AV Block | 57 | 3.4% |  |
| Total patients: 3,070 \| Training: 2,148 \| Validation: 461 \| Test: 461 \| Features: 17 |  |  | Total patients: 1,700 \| Training: 1,190 \| Validation: 255 \| Test: 255 \| Features: 75 |  |  |  |
| Pair | Observed Rate |  | Pair | Observed Rate | Expected Rate | Lift |
| Nephropathy and Neuropathy | 21.8% |  | VT and VF | 0.53% | 0.10% | 5.13 |
| Nephropathy and Retinopathy | 24.7% |  | VT and AV Block | 0.29% | 0.08% | 3.55 |
| Neuropathy and Retinopathy | 24.5% |  | VF and AV Block | 0.35% | 0.14% | 2.52 |
| All three | 11.4% |  |  |  |  |  |

Myocardial infarction (MI) produces a different structure through a different mechanism. Coronary artery occlusion initiates two simultaneous biological processes. The first is body-wide: sympathetic activation floods the myocardium with catecholamines, induces calcium overload, and elevates electrical instability across the entire heart in every patient regardless of which artery is blocked [13]. The second is territory-specific: the infarct location determines which local mechanisms activate. Right coronary artery occlusion threatens the atrioventricular node through direct ischaemia and vagally mediated mechanisms [14]; left ventricular infarcts activate ischaemic border zone reentry mechanisms that generate ventricular tachycardia and drive its degeneration into ventricular fibrillation through Purkinje network fragmentation [15]. The pairwise lift analysis is consistent with this heterogeneous structure: ventricular tachycardia and ventricular fibrillation co-occur at 5.13 times the rate expected by chance, ventricular tachycardia and atrioventricular block at 3.55 times, ventricular fibrillation and atrioventricular block at 2.52 times (Table 1).

Co-occurrence structure is not a single thing. A lift value above one tells you that two complications co-occur more than chance predicts. It does not tell you whether they share a common upstream cause, whether one amplifies the risk of the other, or whether both mechanisms operate simultaneously at different scales. Those distinctions are biologically meaningful and architecturally consequential.

### Inductive Bias as a Biological Hypothesis

Statistical learning theory formalizes what is at stake. The bias-variance tradeoff specifies that a model with appropriate structural constraints reduces variance at the cost of bias, and when the constraint is correct, that trade is beneficial [16, 17]. Theory leaves the correct structural assumption for each problem unresolved.

In multi-label clinical prediction, outcomes may share a common biological cause without directly influencing each other, making that question empirical. A poorly matched inductive bias can hurt performance [18]. We can then ask which structural assumption about comorbidity most accurately reflects the true data-generating process for a given disease, and what does the consistency of its advantage over a well-matched baseline depend on?

Different output-layer architectures encode different biological hypotheses, and the experiment that pits them against each other across data fractions and disease settings is the test of those hypotheses. A symmetric conditional random field makes a specific claim: complications co-occur without directionality, no complication causes the others, and their joint appearance in patients follows from a shared upstream biological cause acting on all of them simultaneously [19, 20]. A multiplicative interaction head makes a different claim: the presence of one complication scales the risk of another through an amplifying relationship, so one outcome multiplies the probability of another proportionally to its own magnitude. A linear additive head claims something different again: that the influence of one complication on another is constant, additive, and independent of the magnitude of the base risk. Each of these is a biological hypothesis stated in architectural form. Choosing between them is a decision about what you believe the disease is doing.

This framing makes the experiment falsifiable in a way that standard benchmarking is not. If the symmetric conditional random field (CRF) wins consistently in a disease where complications share a common upstream cause, that supports the biological claim encoded in the CRF. If the multiplicative head wins in a disease where one complication is known to amplify another, that supports a directed amplifying relationship in the biology. If no single mechanism dominates, that supports the interpretation that the co-occurrence structure is heterogeneous. Every result, clean or mixed, is biologically interpretable. The pattern of which assumption wins may therefore serve as a probe for interaction structure in co-occurrence data—though that reading requires care. A mechanism that wins on AUROC may do so because its structural assumption matches the biology, because its parameter constraints regularise well at a given dataset size, or because the constraint matches dataset noise. Distinguishing those interpretations requires examining whether the advantage is consistent across data fractions and seeds, and whether it aligns with what the biology independently predicts—which is precisely the design the experiment is built around.

The bias-variance argument suggests structural constraints are especially valuable when data is limited. That advantage remains conditional because a simpler regularisation approach—dropout, L2 penalty, or label smoothing—applied to an unconstrained model could achieve comparable variance reduction without any structural assumption at all. That comparison is outside the present study, so the claim that structural grounding helps in data-limited settings should be read with that alternative in view. A constrained model that wins because its assumption is biologically plausible may nonetheless be more interpretable than an unconstrained model that wins for reasons it cannot articulate—because the former is doing something the clinician can recognise, verify, and interrogate.

### What is Missing

The multi-label clinical prediction literature has addressed label dependencies primarily as a learning problem, using chain-based, recurrent, graph-based, and conditional mechanisms to capture co-occurrence from data [6, 19–21]. This data-driven framing leaves less room for disease-specific structural knowledge as an explicit modelling input. The claim here is narrower: clinical complication prediction should choose the form of label interaction from disease biology instead of treating it as a generic dependency-learning problem. In parallel, the biological literature has characterised the comorbidity mechanisms this paper targets in considerable detail: the shared microvascular pathophysiology of diabetic complications [7–9, 12], the sympathetic and ischaemic mechanisms driving electrical complications after myocardial infarction [13, 15], and the territory-specific pathways generating atrioventricular block [14]. What remains underdeveloped is the explicit translation of those disease mechanisms into output-layer assumptions that can be tested against alternatives. This paper treats biological knowledge about comorbidity structure as an architectural input before model training.

This distinction can be written compactly. A standard multi-label neural predictor maps a patient input through a shared representation and an output head,

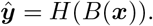

The structured models evaluated here insert an explicit interaction function between the backbone representation and the final predictions,

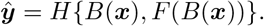

Here, *B*(***x***) is the shared patient representation, *F* encodes an output-layer assumption about how complication labels interact, and *H* maps the representation and interaction signal to predictions. The contribution is the biological choice of *F* : symmetric for shared upstream microvascular damage, multiplicative for amplifying relationships, and unconstrained when no specific structure is assumed. Choosing *F* is therefore a biological claim about the disease, and comparing alternative forms of *F* across data fractions and seeds is the empirical test of that claim.

## Contributions

This paper makes three contributions.

The first is empirical. The T2D setting evaluates six output heads—independent baseline, linear additive, multiplicative, symmetric CRF, residual multilayer perceptron (MLP), and combined additive–multiplicative—across five training data fractions and seven independent seeds. The MI setting evaluates four output heads—independent baseline, linear additive, multiplicative, and symmetric CRF—under the same data-fraction and seed protocol. The central question is whether the accuracy of a structural assumption about comorbidity, grounded in the biology of the disease, determines which mechanism wins and by how much. In T2D the prediction is clean: the symmetric CRF, with three learnable parameters, is expected to show the most consistent advantage over a well-matched baseline across data fractions. In MI the expected pattern is mixed: no single mechanism is predicted to dominate, and the baseline is expected to remain competitive throughout.

The second contribution is interpretive. The pattern of which mechanism wins across data fractions is treated as a biological reading, with performance ranking serving as the evidence for that reading. The identity of the winning mechanism is used as a cautious probe for the biological interaction structure of the disease, especially when the advantage is consistent across data fractions and seeds and aligns with independent biological expectations. The two experiments together map the boundary conditions of that instrument, from the clean symmetric structure of T2D to the heterogeneous structure of MI.

The third is an observation motivating future work. Combining additive and multiplicative interactions without a principled decomposition of which component captures which biological layer does not appear to capture both structures—the combined head shows no consistent advantage and, at lower T2D data fractions, the largest negative deviation from the baseline (Table 3). Architectural combination is coherent only when there is a biological argument for which mechanism encodes which layer of co-occurrence.

The falsifiable claim beneath all three is this: the structural assumption encoded in an output-layer interaction mechanism is a biological hypothesis, and the consistency of its advantage over a well-matched baseline is the test of that hypothesis. A clinical ML practitioner should ask, before choosing an output layer: what does this architecture assume about the disease, and is that assumption defensible?

## Materials and methods

### Datasets

#### Type 2 Diabetes Microvascular Complications

The T2D dataset [22] contains 3,070 clinical records of patients diagnosed with Type 2 diabetes, with 17 input features capturing demographic and clinical variables including age, body mass index, systolic and diastolic blood pressure, glycated haemoglobin, family history, age of onset, smoking status, and medication usage. Dataset characteristics and retained feature counts are summarised in Table 1.

Three binary complication labels were retained as prediction targets: diabetic nephropathy, diabetic neuropathy, and diabetic retinopathy. These three complications share a single generating mechanism—systemic microvascular damage driven by chronic hyperglycaemia—which grounds a specific expectation about the form of their co-occurrence: symmetric, with no directed relationship between specific pairs.

Individual label prevalences were 47.8% for nephropathy (1,468 positive cases), 48.6% for neuropathy (1,492 positive cases), and 50.4% for retinopathy (1,547 positive cases). Pairwise co-occurrence rates were 21.8% for nephropathy–neuropathy, 24.7% for nephropathy–retinopathy, and 24.5% for neuropathy–retinopathy, with 11.4% of patients presenting all three simultaneously. Across all three pairs, the range of co-occurrence rates spans 2.9 percentage points—a near-uniformity consistent with a symmetric upstream mechanism acting on all three microvascular targets simultaneously, with little evidence for directed relationships between specific pairs. Label prevalence and co-occurrence summaries are reported in Table 1.

The dataset was split 70:15:15 into training (2,148 samples), validation (461 samples), and test (461 samples) partitions. Stratified splitting was applied where feasible to preserve label distribution across partitions. Split-level sample counts and overall label prevalences are reported in Table 1.

### Myocardial Infarction Electrical Complications

The MI dataset [23] contains 1,700 clinical records of patients admitted with acute myocardial infarction, with 111 input features. Only features available at the time of hospital admission were included, as specified by the dataset authors’ data split protocol; features that become available only during the hospital period were excluded. After preprocessing and removal of features with excessive missingness, 75 input features were retained. After this preprocessing step, the retained feature count is reported with the dataset characteristics in Table 1.

Three binary complication labels were selected as prediction targets: ventricular tachycardia (VT), ventricular fibrillation (VF), and atrioventricular block (AVB). These complications represent distinct but partially overlapping electrophysiological consequences of acute MI. They arise from a combination of body-wide sympathetic activation and territory-specific ischaemic mechanisms—a heterogeneous co-occurrence structure that contrasts directly with the T2D setting, and that grounds the expectation that a symmetric shared-mechanism model will not necessarily be the best-fitting assumption here.

Individual label prevalences were 2.5% for ventricular tachycardia (42 positive cases), 4.2% for ventricular fibrillation (71 positive cases), and 3.4% for atrioventricular block (57 positive cases). At these prevalences, some label combinations had fewer than two samples, precluding stratified splitting. A random split yielded 1,190 training samples, 255 validation samples, and 255 test samples in a 70:15:15 ratio. Label prevalences, split-level counts, and observed pairwise co-occurrence rates are reported in Table 1.

### Co-occurrence Analysis

Co-occurrence structure in each disease setting was characterised using pairwise lift, defined for complications *i* and *j* as:

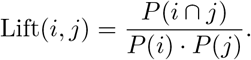

A lift value of 1.0 indicates statistical independence; values above 1.0 indicate co-occurrence more frequent than chance. In the T2D setting, individual prevalences were sufficiently high that raw co-occurrence rates characterised the pairwise structure directly. In the MI setting, individual prevalences below 5% made raw co-occurrence rates near zero and difficult to interpret; lift values were therefore used instead.

In the MI setting, pairwise lift values were 5.13 for ventricular tachycardia and ventricular fibrillation, 3.55 for ventricular tachycardia and atrioventricular block, and 2.52 for ventricular fibrillation and atrioventricular block. The VT–VF pair shows approximately twice the lift of the VF–AVB pair. This asymmetry is consistent with the shared ischaemic border zone substrate and electrophysiological progression pathway that links the ventricular pair—a mechanism atrioventricular block does not participate in to the same degree. This analysis was conducted before model training to establish that interaction structure exists in both datasets and that it differs between settings in a way that carries a biological interpretation. Pairwise co-occurrence and lift summaries are reported numerically in Table 1 and visually in Fig 1.

**Fig 1**. Pairwise co-occurrence structure in the T2D (left) and MI (right) disease settings. In the T2D setting, off-diagonal cells span 2.9 percentage points—the near-uniformity expected from a shared upstream cause acting symmetrically on all three targets. In the MI setting, the VT–VF pair shows lift of 5.13, approximately twice the VF–AVB lift of 2.52, consistent with the distinct electrophysiological substrates that link the ventricular pair and that atrioventricular block does not share. MI diagonal cells are shown as em dashes because self-comparisons are not interpreted as pairwise lift values.

### Shared Backbone Architecture

All models evaluated in this study share a common backbone, following the shared-representation logic of multi-task and multi-label learning [24–28]:

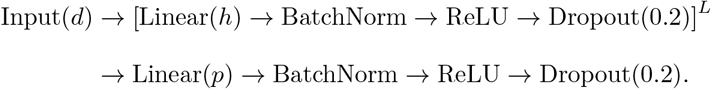

where *d* is the input dimension, *h* the hidden dimension, *L* the number of hidden layers, and *p* the projection dimension. The backbone produces a fixed-dimensional representation ***h*** ∈ ℝ^*p*^, which is passed to the output layer interaction mechanism. Backbone configurations and parameter counts are summarised in Fig 2.

**Fig 2**. Shared feed-forward backbone shown as a traditional node-layer neural network. Input features *x*_1_, …, *x*_*d*_ pass through *L* repeated hidden blocks, each consisting of Linear(*h*), BatchNorm, ReLU, and Dropout(0.2), followed by a final Linear(*p*), BatchNorm, ReLU, and Dropout(0.2) projection. The resulting representation ***h*** ∈ ℝ^*p*^ is passed to every output-layer interaction mechanism.

### Backbone Constraint Rationale

A constrained backbone is what makes the interaction mechanism the experimental variable. A backbone with sufficient capacity to approximate the joint label distribution on its own renders the output layer less informative—any head may perform similarly because the backbone has already encoded much of the co-occurrence structure present in the data. Constraining the backbone to a fixed parameter budget ensures that the output layer is the primary location in the architecture where label interaction can be captured.

### Samples-to-Parameter Ratio

Backbone configurations were selected by samples-to-parameter ratio, targeting a ratio of approximately 2.0; validation performance was not the selection criterion. At this ratio, the model has enough capacity to represent the input structure without enough parameters to compensate easily for a mismatched output layer assumption through memorisation.

For the T2D setting, the backbone was configured with hidden dimension 16, three hidden layers, and projection dimension 8, applied to an input of dimension 17. This yields approximately 1,107 learnable parameters and a ratio of 1.94 at full training data. For the MI setting, with input dimension 75 and a smaller training set, the backbone was reconfigured to hidden dimension 6, one hidden layer, and projection dimension 8, yielding 567 parameters and a ratio of 2.10. Despite different input dimensions, dataset sizes, and absolute parameter counts, the two settings operated at comparable ratios. The backbone settings and parameter counts are summarised in Fig 2; the corresponding samples-to-parameter ratios are reported here to document the matching criterion.

### Depth Sweep

A depth sweep was conducted across backbone configurations with hidden layers from 1 to 8 at hidden dimension 128, evaluated on the T2D dataset at full training data. At each depth, both the baseline and the linear additive head were evaluated on validation macro AUROC, and the gap between them was recorded. The gap as a function of backbone depth characterises how backbone capacity affects the interaction mechanism’s contribution—as depth increases and the backbone absorbs more of the joint label structure, the gap is expected to close. The configurations that sustain a meaningful gap informed the search range for the samples-to-parameter criterion described above; full sweep results are reported in Table S2.

### Baseline Model

The baseline appends an independent linear classification head to the backbone:

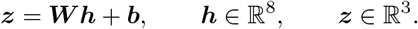

Each logit is computed independently, and binary cross-entropy loss is applied across all three outputs jointly. This binary-relevance construction encodes the assumption that the three complication labels are conditionally independent given the input features [3, 4]. It is the null hypothesis: no structural assumption about comorbidity outperforms the absence of one.

### Interaction Mechanisms

All interaction mechanisms receive the same shared-backbone representation and begin from the same independent baseline logits,

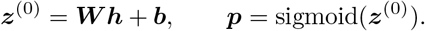

The mechanisms differ only in how they allow the three predicted complication probabilities to modify one another before the final output. The T2D task evaluated the full set of heads: independent baseline, linear additive, multiplicative, symmetric CRF, residual MLP, and combined additive–multiplicative. The MI task evaluated the independent baseline, linear additive, multiplicative, and symmetric CRF heads. The residual MLP and combined head were restricted to T2D, where the full parameter-efficiency and mechanism-combination questions were tested.

**Fig 3**. Output layer interaction mechanisms evaluated in this study. Each mechanism receives the same backbone representation ***h***, base logits ***z***^(0)^, and base probabilities ***p***; only the output-layer interaction assumption changes. Parameter counts are additional to the shared baseline head. Cards marked T2D + MI were evaluated in both disease settings, whereas the residual MLP and combined head were evaluated in the T2D setting only.

### Linear Additive and Multiplicative Heads

The linear additive head adds a masked weighted sum of the other labels’ probabilities to each base logit:

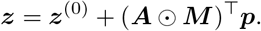

The multiplicative head instead scales each base logit by a learned cross-label factor:

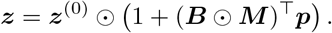

Here ***A*** and ***B*** are 3 × 3 learnable interaction matrices and ***M*** is a fixed zero-diagonal mask that prevents self-interaction. Each head adds six off-diagonal interaction parameters. The additive form represents a constant cross-label contribution, whereas the multiplicative form represents risk amplification proportional to the base logit [4, 6, 29, 30].

### Symmetric Conditional Random Field

The symmetric CRF represents undirected pairwise dependence among labels. Only the three upper-triangular pairwise terms are learned and mirrored to form a symmetric interaction matrix ***β***. Mean-field inference is initialised at the base probabilities and run for three iterations [19, 20, 31–33]:

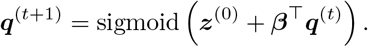

Final logits are recovered as log{***q****/*(1 ***q***)}. This head is biologically aligned with T2D, where the complication structure is expected to be symmetric through shared microvascular damage, and serves as a direct test of whether that same symmetric assumption is too restrictive for MI.

### Residual Multilayer Perceptron

The residual MLP is the unconstrained comparator. It learns an interaction correction from the concatenated backbone representation and base probabilities:

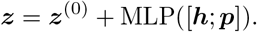

The MLP has one hidden layer of 8 units and a zero-initialised output layer, so training begins from the independent baseline. With 123 additional interaction parameters, it tests whether extra flexible capacity can outperform a smaller biologically matched mechanism [34].

### Combined Additive–Multiplicative Head

The combined additive–multiplicative head is introduced here to test whether two interaction forms can be learned jointly without first assigning each component a distinct biological role. It computes separate additive and multiplicative transformations and then combines them:

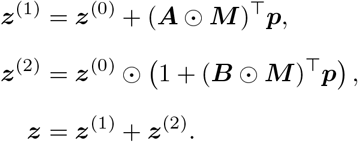

where ***A*** and ***B*** are separate masked 3 × 3 interaction matrices. In simplified form, the head can be understood as

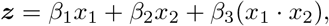

with independent additive, amplifying, and joint-effect terms. Twelve learnable interaction parameters are added—six additive and six multiplicative. This head therefore tests whether combining mechanisms improves robustness or whether the two components compete for the same limited co-occurrence signal.

### Shared Initialisation Protocol

Within each experimental condition defined by a seed and data fraction, all models shared an identical backbone initialisation. At the start of each condition, the backbone was instantiated and its state was saved. Before each interaction model was trained within that condition, the backbone was reset to this saved state. Differences in output-layer performance across models within a condition cannot, therefore, be attributed to different initialisation points.

### Training Protocol

All models were trained using Adam with learning rate 1 × 10^−3^ and batch size 32 [35]. Binary cross-entropy with logits loss was applied independently to each label and summed across labels at each training step. Models were trained for a maximum of 300 epochs with early stopping on validation macro AUROC, retaining the weights at peak validation performance. The same protocol was applied across all models, interaction mechanisms, and disease settings.

### Experimental Design—Data Fraction Analysis

Each model was evaluated across five training data fractions: 100%, 80%, 60%, 40%, and 20% of the available training set. At each fraction, the subset was drawn by sampling without replacement from the full training set using a fixed random state determined by the experimental seed, ensuring that smaller fractions are strict subsets of larger ones within the same seed.

The central experimental question is whether the advantage of a structurally correct output-layer assumption grows, shrinks, or remains stable as training data becomes scarce. Five fractions spanning near-minimal to full data were selected to answer that question while remaining computationally tractable across all models, seeds, and disease settings (Table 2).

**Table 2.** Data-fraction experimental design. Each row represents one training-data fraction evaluated across seven random seeds. The T2D setting uses six output heads: baseline, linear additive, multiplicative, symmetric CRF, residual MLP, and combined additive–multiplicative. The MI setting uses four output heads: baseline, linear additive, multiplicative, and symmetric CRF. Smaller fractions are strict subsets of larger fractions within each seed.

| Training fraction | T2D runs | MI runs | Nested sampling relation |
| --- | --- | --- | --- |
| 20% | 7 seeds × 6 heads | 7 seeds × 4 heads | Smallest subset |
| 40% | 7 seeds × 6 heads | 7 seeds × 4 heads | Strict subset of 60% |
| 60% | 7 seeds × 6 heads | 7 seeds × 4 heads | Strict subset of 80% |
| 80% | 7 seeds × 6 heads | 7 seeds × 4 heads | Strict subset of 100% |
| 100% | 7 seeds × 6 heads | 7 seeds × 4 heads | Full training set |
| Total | 5 fractions × 7 seeds × (6 T2D heads + 4 MI heads) = 350 training runs | | |

### Variance Estimation

Each experiment was replicated across seven independent random seeds: 42, 123, 456, 789, 1337, 2024, and 9999. Seeds controlled both the subsampling of training data at reduced fractions and the initialisation of model parameters. Results are reported as mean macro AUROC ± one standard deviation across seven seeds at each data fraction. At the dataset sizes involved, seven seeds are sufficient to distinguish consistent directional advantages from seed-sensitive fluctuations while remaining computationally feasible across the full set of experimental conditions.

### Evaluation Metrics

The primary evaluation metric was macro-averaged AUROC across the three complication labels. Macro averaging assigns equal weight to each label regardless of prevalence—in the MI setting, where individual complication prevalences fall below 5%, this prevents rarer labels from being discounted in aggregate performance estimates. AUROC evaluates ranking performance across the full range of decision thresholds, providing a more complete characterisation of model discrimination than threshold-dependent metrics at a fixed operating point [36].

Macro F1 score, Hamming loss, and subset accuracy were computed as secondary diagnostics, but the experimental comparisons and conclusions are based on macro AUROC [3, 4].

### Implementation

All models were implemented in PyTorch [37]. Experiments were conducted in Google Colab. Random seeds were set explicitly across Python, NumPy, and PyTorch at the start of each experimental condition. Backbone initialisations were saved as state dictionaries and reloaded before each model was trained within a condition. Results for each seed-fraction condition were written to structured JSON files and copied to Google Drive upon completion. Full per-seed, per-fraction results are available in the supplementary materials.

## Results

### Co-occurrence Structure

The co-occurrence structure of each disease setting was characterised before model training to establish what form of label interaction the output layer should be designed to capture (Table 1 and Fig 1). In the T2D setting, pairwise co-occurrence rates span 21.8% to 24.7% across all three complication pairs—a 2.9 percentage point range across all three pairs. This near-uniformity is the signature of a symmetric upstream mechanism acting on all three microvascular targets simultaneously, with little evidence for directed relationships between specific pairs. In the MI setting, individual complication prevalences of 2.5–4.2% make raw co-occurrence rates uninformative; lift values carry the signal. The VT–VF pair shows lift of 5.13; the VF–AVB pair, 2.52—a 2:1 ratio between the two extreme pairs. This asymmetry is consistent with the shared ischaemic border zone substrate linking the ventricular pair, a mechanism atrioventricular block does not participate in to the same degree. These two settings were selected because they ground different structural expectations at the output layer: symmetric in T2D, heterogeneous in MI.

### T2D: Symmetric Prior Versus Unconstrained Learning

In the T2D setting, the symmetric CRF achieved the highest mean AUROC at full data and at the two lowest training fractions in the full all-model aggregate comparison (Table 3). This aggregate table answers which model has the highest mean AUROC among all mechanisms at each fraction. The paired CRF-versus-baseline comparison answers a different question: whether the CRF improves over the same baseline under matched seed-fraction conditions. In that paired comparison, the CRF is higher than the baseline at every data fraction, with the largest advantage at 20% training data (Table 4, Fig 4). The combined head’s deviation from the baseline became strongly negative once data were reduced to 60% and remained the largest negative deviation of any mechanism at 60%, 40%, and 20% training data, consistent with gradient interference between the additive and multiplicative components becoming more pronounced under data scarcity. The structural assumption, not the parameter count alone, determines which mechanism holds an advantage.

**Table 3.** Mean macro AUROC across seven seeds—Type 2 diabetes microvascular complications. Values are reported as mean (standard deviation) across seven independent random seeds. Bold indicates the highest mean AUROC at each fraction. CRF interaction parameters: 3. Residual MLP interaction parameters: 123. At the two lowest data fractions, the symmetric CRF has the highest mean AUROC; at 20% training data, it is the only interaction mechanism above the baseline. At 60% training data and below, the combined head shows the largest negative deviation from the baseline of any mechanism evaluated.

| Fraction | Train N | Baseline | Linear Additive | Multiplicative | CRF | Residual MLP | Combined Head |
| --- | --- | --- | --- | --- | --- | --- | --- |
| 100% | 2,148 | 0.6321 (0.0135) | 0.6381 (0.0218) | 0.6313 (0.0212) | **0.6430 (0.0248)** | 0.6237 (0.0204) | 0.6316 (0.0223) |
| 80% | 1,718 | **0.6350 (0.0202)** | 0.6290 (0.0306) | 0.6250 (0.0110) | 0.6225 (0.0089) | 0.6302 (0.0165) | 0.6325 (0.0190) |
| 60% | 1,288 | **0.6196 (0.0053)** | 0.6066 (0.0076) | 0.6046 (0.0209) | 0.6099 (0.0215) | 0.6085 (0.0177) | 0.5939 (0.0237) |
| 40% | 859 | 0.5916 (0.0073) | 0.5942 (0.0117) | 0.5853 (0.0199) | **0.5945 (0.0214)** | 0.5819 (0.0129) | 0.5753 (0.0290) |
| 20% | 429 | 0.5648 (0.0257) | 0.5583 (0.0189) | 0.5520 (0.0184) | **0.5675 (0.0225)** | 0.5603 (0.0188) | 0.5506 (0.0182) |

**Table 4.** Paired CRF-versus-baseline comparison across training fractions. Differences are computed within matched seed-fraction conditions before averaging, so the column tests whether the symmetric CRF improves over the same baseline initialisation and data split. This paired comparison is distinct from the all-model aggregate comparison in Table 3. The CRF is higher than the baseline at every fraction, with the largest paired advantage at 20% training data.

| Fraction | Train N | Baseline AUROC | CRF AUROC | $\Delta$ CRF–Baseline | Paired Winner |
| --- | --- | --- | --- | --- | --- |
| 100% | 2,148 | $0.6322 \pm 0.0224$ | $0.6430 \pm 0.0248$ | $+0.0109 \pm 0.0295$ | **CRF** |
| 80% | 1,718 | $0.6184 \pm 0.0138$ | $0.6225 \pm 0.0089$ | $+0.0040 \pm 0.0175$ | **CRF** |
| 60% | 1,288 | $0.6056 \pm 0.0193$ | $0.6099 \pm 0.0215$ | $+0.0043 \pm 0.0180$ | **CRF** |
| 40% | 859 | $0.5808 \pm 0.0166$ | $0.5845 \pm 0.0214$ | $+0.0036 \pm 0.0126$ | **CRF** |
| 20% | 429 | $0.5499 \pm 0.0185$ | $0.5675 \pm 0.0225$ | $+0.0176 \pm 0.0114$ | **CRF** |

**Fig 4**. Paired CRF-versus-baseline performance across training fractions. The left panel shows the paired baseline and CRF AUROC values from Table 4; the right panel plots the paired CRF-minus-baseline difference. The CRF advantage remains positive at every fraction and is largest at 20% training data. Error bars show one standard deviation across seven seeds.

**Fig 5**. Model performance across training data fractions—Type 2 diabetes microvascular complications. At 20% training data, the symmetric CRF is the only interaction mechanism above the baseline; all other interaction mechanisms show negative mean differences at that fraction. The combined head’s negative deviation from the baseline is largest at 60%, 40%, and 20% training data, consistent with gradient interference between the additive and multiplicative components becoming more pronounced under data scarcity.

**Fig 6**. Symmetric CRF versus residual MLP—parameter efficiency across data fractions. Differences are computed from the reported mean AUROC values in Table 3. At 20% and 40% training data, the symmetric CRF maintains a positive mean difference from the baseline, while the residual MLP, with 41 times more interaction parameters, falls below the baseline at every fraction. Three parameters constrained by a biologically grounded structural assumption outperform 123 unconstrained interaction parameters at the lowest data fractions.

### MI: Heterogeneous Structure and Non-Monotonic Competition

The MI results are a finding about biological heterogeneity, not a failure of any single mechanism. The winner changes at three of the four fraction transitions from 20% to 100%—a pattern that is the expected signature of a disease combining body-wide sympathetic activation with territory-specific ischaemic amplification between the ventricular pair. At 100% training data, the CRF leads all three interaction mechanisms, achieving 0.7752 against a baseline of 0.7632. At 80% and 60%, the multiplicative head leads, with mean AUROC values of 0.7873 and 0.7507 respectively. At 40%, the baseline leads all interaction mechanisms at 0.6697, consistent with co-occurrence signal being insufficient at that sample size for any structural assumption to establish a consistent advantage. At 20%, the CRF recovers the lead, achieving 0.6040 against a baseline of 0.5886. The clean separation observed in the T2D paired CRF comparison does not appear here; every line crosses at least one other between fractions, and no mechanism holds its position across the full range.

**Fig 7**. Model performance across training data fractions—myocardial infarction electrical complications. At 100% training data, the CRF leads by 0.0120 AUROC over the baseline; at 60–80%, the multiplicative head leads; at 40%, the baseline leads all three interaction mechanisms. The winner changes at three of the four fraction transitions. The crossing pattern is the expected signature of a disease that combines body-wide sympathetic activation—which a symmetric CRF can exploit at sufficient sample sizes—with territory-specific ischaemic amplification between the ventricular pair, which the multiplicative head captures at medium fractions.

**Fig 8**. Baseline-relative aggregate performance across output heads and training fractions. Cells show mean macro AUROC for each interaction mechanism minus the independent-head baseline mean at the same training fraction, computed from Table 3 and Table 5. Positive values indicate performance above the baseline; negative values indicate performance below the baseline. These are aggregate mean differences, not paired seed-wise differences; the paired T2D CRF comparison is reported separately in Table 4 and Fig 4.

**Table 5.** Mean macro AUROC across seven seeds—myocardial infarction electrical complications. Values are reported as mean (standard deviation) across seven independent random seeds. Bold indicates the highest mean AUROC at each fraction. At 100% training data, the CRF leads all three interaction mechanisms, achieving 0.7752 against a baseline of 0.7632. At 60–80%, the multiplicative head leads. At 40%, the baseline leads all interaction mechanisms; at this sample size, co-occurrence signal is insufficient for any structural assumption to provide consistent benefit. At 20%, the CRF recovers the lead. The winner pattern changes at three of the four fraction transitions—a non-monotonic result that reflects a disease mechanism too heterogeneous to be captured by any single structural assumption across the full data range. Standard deviations above 0.10 at several cells follow directly from the 2.5–4.2% individual complication prevalences: at these rates, small shifts in positive case counts across seeds produce large AUROC swings.

| Fraction | Train N | Baseline | Linear Additive | Multiplicative | CRF |
| --- | --- | --- | --- | --- | --- |
| 100% | 1,190 | 0.7632 (0.0206) | 0.7280 (0.1006) | 0.7569 (0.0595) | **0.7752 (0.0341)** |
| 80% | 952 | 0.7287 (0.0627) | 0.7674 (0.0438) | **0.7873 (0.0337)** | 0.7136 (0.1056) |
| 60% | 714 | 0.7185 (0.1164) | 0.7322 (0.0580) | **0.7507 (0.0645)** | 0.6835 (0.1156) |
| 40% | 476 | **0.6697 (0.0771)** | 0.6453 (0.1723) | 0.6446 (0.1373) | 0.6490 (0.1366) |
| 20% | 238 | 0.5886 (0.1487) | 0.5929 (0.1639) | 0.5804 (0.1439) | **0.6040 (0.1459)** |

## Discussion

### Output Layers Encode Biological Assumptions

The results support the central premise of this study: output-layer design is not a neutral implementation detail in multi-label clinical prediction. Each interaction mechanism encodes a claim about how complications relate, and the usefulness of that claim depends on whether it matches the disease process generating the labels. In T2D, where the biology predicts stable, symmetric co-occurrence through shared microvascular damage [7–9, 12], the symmetric CRF showed the clearest and most consistent advantage (Table 3, Table 4). In MI, where electrical complications arise from both shared sympathetic instability and territory-specific ischaemic mechanisms [13–15], no single output-layer assumption dominated across data fractions (Table 5, Fig 7). That contrast is the main interpretive result.

This does not mean that the winning model reveals disease mechanism on its own. A model may perform well because its structural assumption matches the biology, because it regularises effectively at a given sample size, or because it matches the noise structure of the dataset. The biological reading is strongest only when the performance pattern is consistent across data fractions and seeds and agrees with independently known pathophysiology. The value of the experiment lies in the alignment between model advantage and the structural expectations of the two disease settings (Table 6).

**Table 6.** Synthesis of biological structure, architectural expectation, and observed model behaviour. Pairwise co-occurrence and lift values motivate label-dependence expectations but are not interpreted as mechanistic effect-size estimates.

| Field | Synthesis |
| --- | --- |
| **Type 2 diabetes microvascular complications** |  |
| Biological structure | Shared systemic microvascular injury driven by chronic hyperglycaemia; pairwise co-occurrence rates are near-uniform across nephropathy, neuropathy, and retinopathy. |
| Expected best head | Symmetric CRF, because it constrains the output layer to learn undirected pairwise dependence with only three interaction parameters. |
| Observed pattern | CRF has the highest aggregate mean AUROC at full data and at the two lowest data fractions; in the paired comparison, CRF is above the same baseline at every fraction. |
| Interpretation | The result supports the usefulness of a constrained symmetric dependency when the label-dependence signal is relatively uniform. It does not make the CRF weights biological effect-size estimates. |
| **Myocardial infarction electrical complications** |  |
| Biological structure | Heterogeneous structure combining body-wide sympathetic instability with territory-specific ischaemic mechanisms; lift is stronger for VT–VF than for pairs involving AV block. |
| Expected best head | No single simple head is expected to dominate; multiplicative interactions may help when pair-specific amplification is informative, while CRF may help when shared instability dominates. |
| Observed pattern | Winner changes across fractions: CRF leads at 100% and 20%, multiplicative leads at 80% and 60%, and the baseline leads at 40%. |
| Interpretation | The mixed pattern is consistent with heterogeneous label dependence and low-prevalence instability. It supports a cautious interpretation and avoids claiming that one architecture has recovered the underlying biology. |

### Symmetric Structure in Type 2 Diabetes

The T2D results are the clearest evidence that a biologically matched output-layer constraint can be useful. The symmetric CRF achieved its largest paired advantage over the baseline at 20% training data, where the mean difference was +0.0176, compared with +0.0109 at full data (Table 4). Six of seven seeds showed a positive CRF-over-baseline difference at 20%, compared with four of seven at 100% (Table S1). This pattern is consistent with the expected behaviour of a useful structural prior: the constraint helps most when the data are least able to estimate the interaction structure on their own.

The biological reason is straightforward. Nephropathy, neuropathy, and retinopathy are not expected to cause one another directly. They co-occur because chronic hyperglycaemia damages multiple microvascular beds through a shared systemic process [7–11]. A symmetric interaction model therefore encodes the right kind of relationship: undirected, pairwise, and shared across the population. The fact that this structure is represented with only three learnable interaction parameters strengthens the interpretation. The CRF does not win by being more expressive than the alternatives; it wins by being more specific.

The residual MLP provides the strongest contrast. With 123 interaction parameters, it had far greater capacity than the CRF, but it remained below the baseline at every T2D data fraction (Table 3, Fig 6). This suggests that simply giving the model more freedom to learn label interaction is not enough. In this setting, unconstrained interaction learning appears less useful than a smaller mechanism whose constraint matches the biology.

### Heterogeneous Structure in Myocardial Infarction

The MI results show the opposite pattern. The winning mechanism changes across data fractions: the CRF leads at 100% and 20%, the multiplicative head leads at 80% and 60%, and the baseline leads at 40% (Table 5, Fig 7). Taken only as a ranking, this instability is difficult to interpret. Taken in relation to MI biology, it is informative. Electrical complications after MI are generated by more than one mechanism: a shared sympathetic response increases global electrical instability, while territory-specific ischaemia creates local substrates for ventricular tachycardia, ventricular fibrillation, and atrioventricular block [13–15]. No single output-layer assumption captures all of that structure.

The large standard deviations in the MI table reinforce this interpretation. With individual complication prevalences between 2.5% and 4.2%, small changes in which positive cases appear across seeds can produce large AUROC shifts (Table 1, Table 5). For that reason, individual MI cells should not be overinterpreted. The more reliable finding is the absence of convergence on a single mechanism. In contrast to T2D, where one biologically plausible constraint is consistently useful, MI appears to require either a more decomposed architecture or substantially more data before a stable interaction pattern can be learned.

### Parameter Efficiency in Data-Limited Settings

The parameter-efficiency result matters because clinical complication prediction is often data-limited. Many complications are rare, and increasing the number of positive cases requires large registries or long collection periods. Under those conditions, the ability of a small structured head to improve performance at low data fractions is practically important. In the T2D setting, the CRF remained above the baseline across fractions in the paired comparison, while the larger residual MLP did not exceed the baseline in the aggregate means (Table 4, Fig 6). This suggests that a well-chosen structural constraint can be more useful than additional interaction capacity when the available data are limited.

This result does not prove that biological structure is always superior to generic regularisation. Dropout, L2 penalties, label smoothing, or other regularisation strategies applied to an unconstrained model could reduce variance without encoding a disease-specific assumption, consistent with the broader bias-variance framing of structural constraint [16–18]. That comparison was not performed here. The more cautious conclusion is that biologically grounded structure offers an interpretable form of regularisation: when it helps, the benefit can be related back to a clinical mechanism as well as to variance reduction.

### The Limits of Unprincipled Combination

The combined additive–multiplicative head shows that adding more interaction mechanisms does not automatically produce a better biological model. In T2D, the combined head showed no consistent advantage and became most negative relative to the baseline at lower data fractions (Table 3, Fig 8). This is consistent with the idea that two unconstrained interaction components can compete for the same limited signal instead of dividing the biological problem cleanly between them.

Mixed mechanisms may still be appropriate. In fact, MI is exactly the kind of disease setting where a mixed mechanism may be needed, because the biology contains both shared-cause and amplifying components. The failure of the combined head instead clarifies what such a mechanism would require. A future mixed architecture should assign distinct biological roles to its components before training: one branch for symmetric shared-cause dependence, analogous to structured-output dependency models [19, 20], another for directed amplification, analogous to multiplicative or gated interaction mechanisms [29, 30], and a parameterisation or loss structure that prevents the two from redundantly modelling the same signal. Combination is useful only when it is decomposed.

## Conclusion

The central conclusion is that output-layer architecture should be treated as a biological hypothesis in multi-label clinical prediction. When the structural assumption matched the disease process, as in the symmetric CRF for T2D, a small constrained model outperformed larger and less constrained alternatives. When the disease process was heterogeneous, as in MI, no single assumption dominated and the baseline remained competitive. The choice of output layer therefore makes a claim about how the disease generates complications. That claim should be stated, justified biologically, and tested empirically as part of the modelling argument.

## Limitations

This study has several limitations. The backbone was fixed by design to isolate output-layer effects, but a more expressive backbone could absorb co-occurrence structure before the output layer and reduce the apparent advantage of structured heads. The analysis was limited to two disease settings, and the parameter-efficiency result has not yet been tested in registry-scale datasets or external validation cohorts. The MI results should also be interpreted cautiously because low complication prevalence produced large seed-level variance. In addition, the mean-field CRF used a fixed number of inference iterations, and approximation quality was not evaluated separately. Finally, biologically grounded structure was not compared against matched generic regularisation in unconstrained models, leaving open how much of the observed benefit reflects disease-specific structure and how much reflects variance reduction alone.

## Supporting information

**Table S1.** Per-seed CRF performance relative to baseline—Type 2 diabetes. Positive values indicate CRF outperforms baseline for that seed and fraction. At 20% training data, six of seven seeds show a positive CRF-over-baseline difference—the most directionally consistent fraction in the table. At 100%, four of seven seeds show a positive difference; Seeds 1337 and 2024 are persistently negative at higher fractions before reversing at 40% and 20% respectively. The mean CRF advantage is positive at every fraction, and the per-seed variance does not reverse the direction of the aggregate result.

| Fraction | Seed 42 | Seed 123 | Seed 456 | Seed 789 | Seed 1337 | Seed 2024 | Seed 9999 |
| --- | --- | --- | --- | --- | --- | --- | --- |
| 100% | +0.0310 | −0.0181 | +0.0317 | +0.0074 | −0.0328 | −0.0021 | +0.0591 |
| 80% | +0.0370 | +0.0140 | −0.0112 | −0.0170 | +0.0151 | −0.0050 | −0.0047 |
| 60% | +0.0180 | −0.0048 | +0.0390 | −0.0055 | −0.0124 | −0.0152 | +0.0108 |
| 40% | −0.0093 | +0.0210 | −0.0023 | +0.0111 | +0.0024 | +0.0178 | −0.0151 |
| 20% | +0.0013 | +0.0116 | +0.0255 | +0.0063 | +0.0188 | +0.0228 | +0.0372 |

**Table S2.** Backbone depth sweep—baseline validation AUROC and linear additive gap. Sweep conducted at hidden dimension 128 on 100% T2D training data. At depths 1 and 5, the gap between the baseline and the linear additive head is negative, indicating the backbone has already encoded label interaction. At depths 2–4 and 6–8, the gap is positive, ranging from +0.0009 to +0.0143. The depth showing the largest gap (depth 4, +0.0143) was not the selection criterion. The final backbone—hidden dimension 16, three hidden layers, projection dimension 8—was selected by samples-to-parameter ratio: the configuration that reliably leaves label interaction to the output layer by limiting backbone capacity.

| Depth | Baseline Val AUROC | Linear Additive Val AUROC | Gap | Notes |
| --- | --- | --- | --- | --- |
| 1 | 0.9748 | 0.9700 | −0.0053 | Gap negative |
| 2 | 0.9757 | 0.9858 | +0.0056 | — |
| 3 | 0.9870 | 0.9842 | +0.0026 | — |
| 4 | 0.9776 | 0.9884 | +0.0143 | — |
| 5 | 0.9813 | 0.9835 | −0.0049 | Gap negative |
| 6 | 0.9822 | 0.9853 | +0.0113 | — |
| 7 | 0.9818 | 0.9813 | +0.0009 | Near zero |
| 8 | 0.9796 | 0.9841 | +0.0064 | — |

## Acknowledgments

The authors thank the Department of Biomedical Engineering, University of Ghana, for institutional support. The authors also thank Prof. Rama Ramakrishnan of MIT Sloan School of Management for helpful early correspondence on inductive bias, data efficiency, and explicit comorbidity-aware modelling, which helped motivate the framing of this work.

